# Ecological Drivers of Community Cohesion

**DOI:** 10.1101/2022.05.15.491981

**Authors:** Chaitanya S. Gokhale, Mariana Velasque, Jai A. Denton

**Affiliations:** Research Group for Theoretical Models of Eco-evolutionary Dynamics, Department of Evolutionary Theory, Max Planck Institute for Evolutionary Biology, August-Thienemann-Straße 2, 24306 Plön, Germany; Genomics & Regulatory Systems Unit, Okinawa Institute of Science & Technology, Onna-son, 904-0404, Japan; Experimental Evolutionary Biology Lab, Monash University, Clayton, 3800, Australia; World Mosquito Program, Institute of Vector-borne Disease, Monash University, Clayton, 3800, Australia

**Keywords:** mutualism, evolutionary dynamics, ecological processes, persistence, cycling

## Abstract

From protocellular to societal, networks of living systems are complex and multi-scale. The assembly of these intricate interdependencies, under ecological pressures, can be nearly impossible to understand using pairwise methods. We develop a mathematical and computational model based on a four-strain *Saccharomyces cerevisiae* synthetic inter-dependent system. Our system leverages transiently structured ecologies for achieving community cohesion. We show how ecological interventions could reverse or slow the extinction rate of a cohesive community. An interconnected system first needs to persist long enough to be a subject of natural selection. Our emulation of Darwin’s warm little ponds’ with an ecology governed by transient compartmentalisation provides the necessary persistence. Our results find utility across scales of organisation, stressing the importance of cyclic processes in major evolutionary transitions engineering of synthetic microbial consortia and conservation biology.

## Introduction

Complex living systems are composed of a web of interactions. These interactions are usually structured both in space (i.e. alongside an environmental gradient or conditions) and in time. Temporal and spatial modifications in the environment can modify both the direction and strength of interactions. For instance, tropical savanna is characterised by highly seasonal rainfall, with pronounced dry seasons of varying duration. The high irradiance and heat present during dry seasons can create a water deficit causing dramatic changes in vegetation structure and associated biodiversity, resulting in widespread mortality [1, 2]. Similarly, periods with high rainfall will increase resource availability, leading to an increase in vegetative growth, increasing species diversity, and increasing the interactive-potential [3]. Although rainfall has a strong and somewhat predictable effect on savannah’s diversity and abundance, this is not always the case and can be associated with a decrease in species abundance [4]. In this study, we study how communities of replicators in the most general sense, from prebiotic molecules to species, can persist in such dynamic ecological processes. We use the simplicity of minimal synthetic systems and corresponding theoretical models to achieve our aim.

The complexity of dynamic ecologies can significantly obstruct understanding of community properties such as temporal and spatial components, stability, and even the type of interaction that connects nodes (antagonistic, mutualistic, trophic and such). The loss of a single node can cascade through the network in unpredictable ways. One example is that of the “living dead” [5, 6]. Although still alive in the environment, a node can lack the ecological circumstances that allow them to be reproductive. In some cases, such circumstances can be reversed with intervention (i.e. reforestation, construction of ecological corridors) [7], however, extinctions of mutualist partners typically lead to co-extinction (mutualism coextinction) [8–10], as observed in trees that lost their pollinators [11]. Further difficulties arise when interactions are dynamic, causing a significant indirect effect that can modify species diversity in a non-quantifiable manner, such as interference and facilitation [12–15]. Such complex natural systems can significantly impede our understanding of critical environmental components, such as abundance, resilience, diversity, species interactions and other ecological network properties.

To avoid issues associated with indirect, weak and convoluted interactions across longer timescales or environments, it helps partition species interactions into their ecological role using functional groups [16]. Thus node-rich networks can be expressed in terms of functional diversity rather than species. This approach aims to understand how functional groups influence diversity and how systems adapt to changes in conditions, including environmental degradation. Further still, functional networks can occupy multiple scales, from molecular interaction to planet-wide biomes. This generality allows exploration of how small changes can ripple throughout a network at one level, either biotic or abiotic.

Unpacking this complexity may seem like a Sisyphean task, but this challenge is what a growing body of interlinked, highly tractable theoretical and experimental models seeks to overcome [17–21]. Despite being reductionist, these approaches elucidate core principles of complex community dynamics through tightly controlled interactions. Synthetic systems, and corresponding theoretical models, allow specific control of biotic and abiotic components, ecological structure, nutrients and growth conditions that allow precise measurement of the interaction outcomes and dynamics [18]. Several synthetic microbial systems and closely linked theoretical models have been developed in this case. They fall into two categories: those with fixed functional diversity [17, 22, 23], and those where the function of individual components is permitted to evolve [20, 21], becoming evolutionarily stable. As these systems continue to improve and expand, we envision systems of greater complexity combining both categories.

We build on this existing work by designing a theoretical model informed by experimental data. While the theory developed herein is inspired by the vast complexity of interactions across scales of organisation, we focus on the ecological factors at the root of incipient communities. We aim to show that even if units of functional dependency exist, ecology plays a conducive role in the eventual assembly of a system of stable interdependencies. This includes displacement of biotic functional groups through fluctuating abiotic factors.

Specifically, we develop a theoretical model with four interacting components informed by our engineered four-strain cross-feeding inter-dependent yeast system. The corresponding theoretical model is based on the classical hypercycle [24]. The model of a hypercycle has been used to describe systems of reactants that catalyse the replication of each other with implications from the origins of life to ecosystems [25, 26]. The experimental and theoretical systems posit that fluctuating ecological processes can assemble functionally dependent but unconnected components into stable communities. This process facilitates exploration of the complex dynamics between species’ functional diversity, environmental context and how their combination shapes the interactions in the lineage that survives. Instead of evolving symbiotic interactions, we start with engineered interactions [17, 22]. This approach allows us to control one aspect of eco-evolutionary dynamics, the interactions, while exploring the ecology [27, 28]. Using the growth properties of this system, we develop a theoretical model. We demonstrate how small ecological changes combined with the community sampling and merging influence system survivorship. Moreover, we show how ecological interventions can stave off system-wide extinction.

## Model & Results

Evolutionary and ecological constraints drive symbiosis in Nature [29]. Hence, our model and subsequent analysis, are divided to reflect the ecological and evolutionary dynamics underpinning functional interdependence. These two core processes are first studied independently and then combined to study the joint eco-evolutionary trajectory.

### Strains

Our system relies on four-way metabolite cross-feeding with one of the four strains, each containing feedback-resistant (FBR) mutation that overproduce either adenine, tryptophan, histidine or lysine. Several FBR mutations in numerous biosynthetic pathways have been identified that facilitate yeast cross-feeding [30]. This has facilitated the development of synthetic yeast cross-feeding models of mutualism. A system of interdependent strains of yeast, one overproducing lysine and lacking adenine while the other overproducing adenine and lacking lysine, was developed by [22] and analysed in detail both biochemically and ecologically [17, 28]. The strains overproducing leucine and lacking tryptophan and vice versa were developed by [23].

Here we developed a four-strain system that relied on the previously referenced FBR mutants. We constructed strains overproducing one of the metabolites while requiring the other three. For example, strain *ADE↑* contains a mutation in ADE4 that prevents the biosynthetic pathway from being inactivated. The resulting construct overproduces adenine and requires tryptophan, leucine, and histidine for growth. Similar mutations were employed in each of the four strains *ADE↑, TRP↑, HIS↑* and *LYS↑* (Materials & Methods). With all the metabolites present *ad libitum*, the growth rates of these engineered strains are calculated (Figure 1).

**Figure 1:**
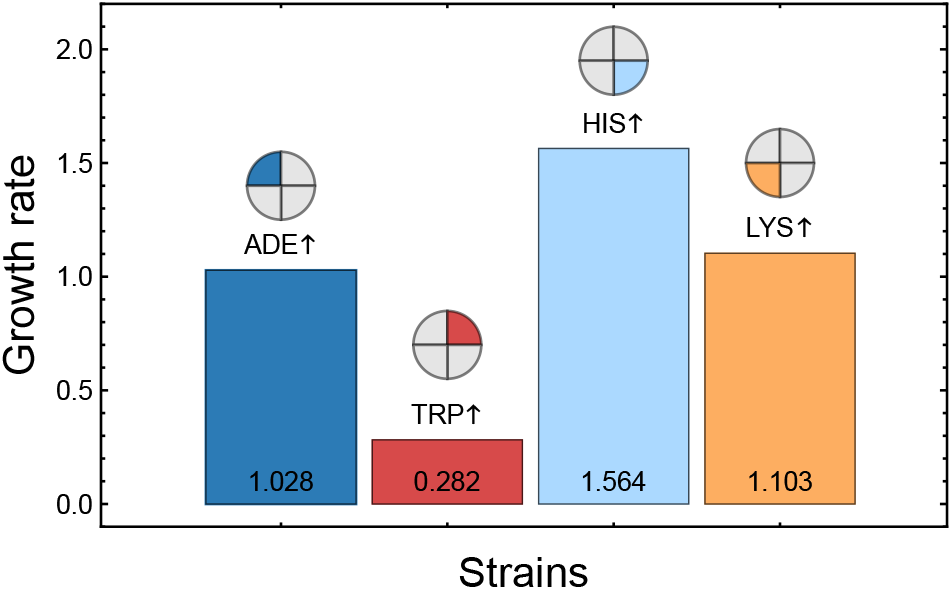
Growth rates of the four engineered strains of the synthetic mutualism system in minimal media. Starved cultures of each of the four strains were diluted in minimal media and pooled. Growth rates were derived from three-day averages in minimal media. All experiments were conducted in triplicate. The minimal media environment (yeast synthetic complete lacking amino acid supplementation) required four-way cross-feeding for strain growth. See Materials & Methods for further details of the experimental setup.

Synthetic mutualism was determined through growth in conditions that required four-way cross-feeding. We calculated growth rates in all pairwise combinations of the engineered strains and with the four pooled strains. From these experiments, we extrapolate growth rates of the individual strains when present in different community compositions and when in a variety of ecologies (see Materials & Methods - Estimating growth parameters).

### Ecological processes

#### Ecologies

The strains are defined by their ability to produce metabolites. Therefore we define the different possible ecologies also by the presence or absence of the four essential metabolites ADE, TRP, HIS and LYS. Thus, in principle, sixteen distinct ecological conditions, from extremely poor - devoid of any metabolite, to rich - where all the metabolites are present, can be visualised Fig. 2. To begin with, we assume that all the sixteen ecologies have the same probability of being realised.

**Figure 2:**
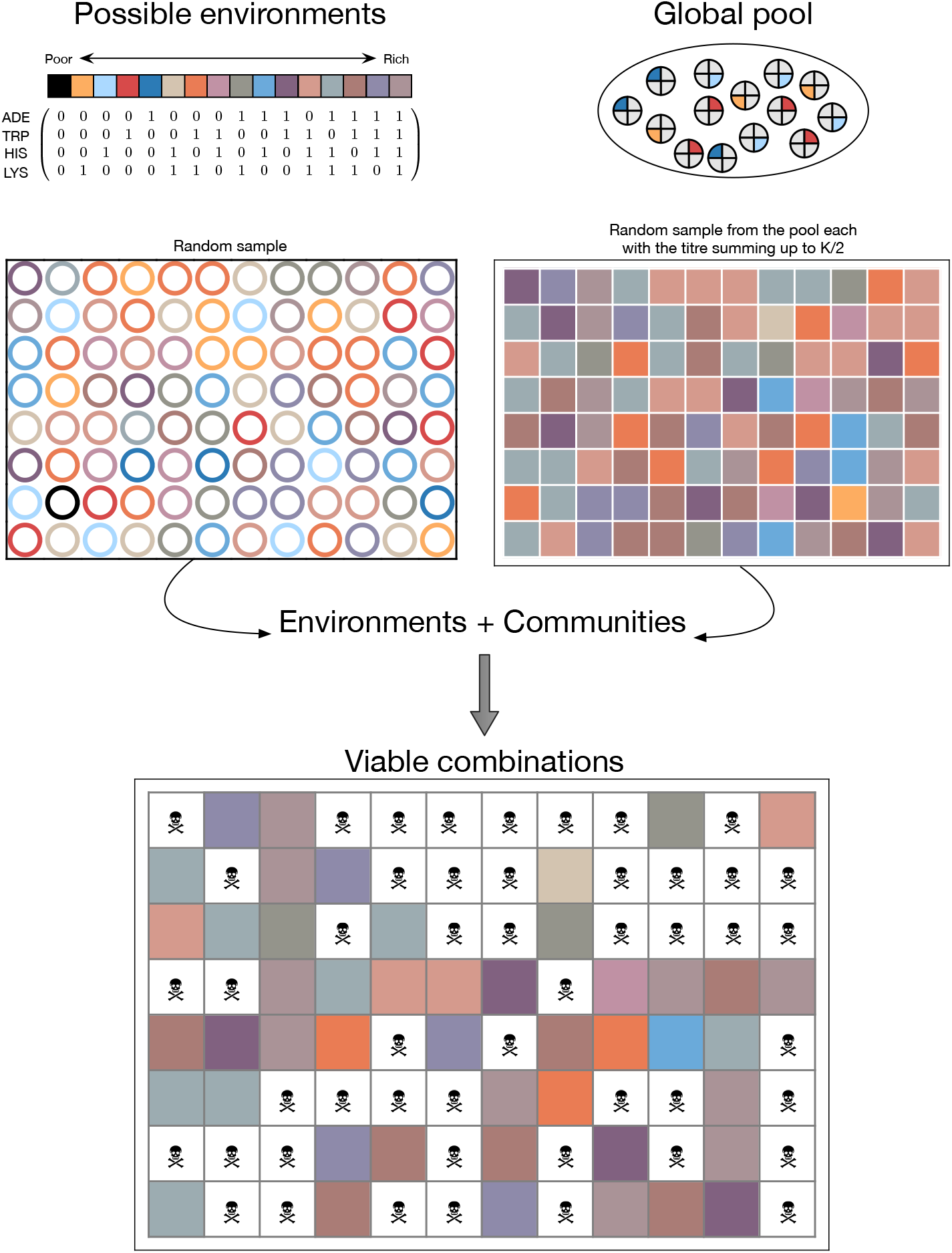
Perfect match. While the possible ecological range from rich (1,1,1,1) to poor (0,0,0,0), the community compositions are a random sample of four types of strains. Given the carrying capacity of each well to be *K*, we sample *K/*2 amount of strain. The resulting communities may be a mixture of all the four strains (*K/*8, *K/*8, *K/*8, *K/*8) or a monoculture e.g. only ADE+ (*K/*2, 0, 0, 0) and all other possible combinations in between. Only when the community members have access to all the required metabolites, either through crossfeeding or the ecology, then the combination is termed as viable. For example in the top-left cells the ecology is (1,0,0,0), i.e. only adenine is present in the ecology and the community consists of Ade+, His+ and Lys+. The strains require tryptophan which is not available and hence the strains will die. Instead, in cell {1,4}, the community composition is the same but the ecology consists of TRP (along with HIS and LYS) and thus is a viable ecology. The difference in growth rate comes from the fact that cross-fed strains have a different growth rate than supplemented (e.g. see cells {6,7} and {6,8}).

Resources can typically vary spatially as well as temporally. For example, metabolites can heterogeneously spread in space with differing amounts. This scenario is reminiscent of the pores in hydrothermal vents where nutrients coalesce for a limited time before being washed out by physical processes. We include this variability in our model by considering different possible ecological conditions. For simplicity, we visualise this diversity in 96 cultures reflecting a 96 well-plate experimental approach Fig. 2. We sample the aforementioned sixteen possible ecological states (here uniformly) to fill the 96 positions.

Next, we inoculate each ecology with a microbial community. Communities of strains are generated by sampling from a global pool of strains Fig. 2. Physical limits of ecologies can support only a limited number of individuals. Hence, the carrying capacity of each of the wells on the plate is limited to *K*. The composition of each community can at most consist of four different strains. This combinatorics leads to 256 distinct possibilities from the pool. For inoculation, the community is scaled to *K/*2, i.e. half the carrying capacity. Thus while group sizes (here proxied by the density *K/*2) are fixed, the compositions are heterogeneous.

#### Within well dynamics

Each of the 96 wells is thus a combination of the ecological state and a community. The required metabolites can be available via cross-feeding or already present in the ecology for each strain in a community. Suppose the requirements of all the strains in each community are satisfied, we categorise them as viable Fig. 2. Within each well the growth of each of the four strains is given by,

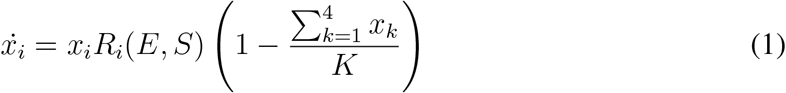

Here, *R*_*i*_(*E, S*) captures the different growth rates depending on the ecology of the well *E* and the strain composition of the community *S*. Each metabolite could be either present or absent in each well. If present, the ecology could be the source of the metabolite, a cross-feeding strain, or both. Growth rates of strain that rely on uptake, due to being unable to produce a specific nutrient, can be adversely impacted [31]. For various such combinations of strains and ecologies, we have inferred the relevant growth rates for each strain coming from minimal experiments in Fig. 1.

All strains are assumed to have the same death rate *d*. If a strain does not have access to all required metabolites, provided through either the community or ecology, it suffers this negative growth rate. In an alternative version of our model we set the *d* = 0 which leads to persistence but this does not affect the qualitative result of our model (code available online). Thus if the ecologies and communities do not match or if the strains themselves are mismatched, then those communities, as a whole, shrink Fig. 2. The concentration phase in the wells lasts for *t*_*wells*_ time-steps.

#### Cycling

We implement an ecological cycle to consider the dynamic nature of fluctuating environmental conditions. The cycle consists of a concentration phase denoted by the 96 wells described above and a pool phase where the strains enjoy a nutrient-rich environment unconstrained by space. At the end of the wells-phase duration, the concentrations are normalised, giving us the input for the next pool phase. The pool is assumed to be nutrient-rich, and the strains have no dearth of space. Hence the growth rates for the four strains are assumed to be the same as represented by the fitnesses in Fig. 1. In the pool the strains grow exponentially,

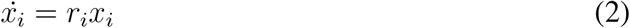

where *r*_*i*_ is the growth rate of strain *i* as given in Fig. 1. This process runs for a time duration *t*_*pool*_. At the end of *t*_*pool*_ we sample the pool to prepare the inoculums for the concentration phase. The ecologies and the communities for each of the 96 wells are prepared as per the procedure describe before.

The result of such an exemplary pool-wells-pool cycle is shown in Fig. 3. Since the ecology and community sampling add a level of randomness to the process, we show the average number of runs in the bottom panels of Fig. 3. The trajectories of the four strains show that the result overall is driven by the pool fitnesses where the *HIS↑* has the highest fitness and thus takes over the population. Since there can be sixteen different ecologies possible, the only ones able to support an all *HIS↑* community are i). lacking in histidine and ii). all metabolites are present, which can occur with a probability of 0.0625 each. Thus the upper bound on the proportion of death that one can expect to see in repeated pool-well cycles over a long time is 0.875. Due to non-zero proportions of *ADE↑* and *LYS↑*, the observed values are lower than expected.

**Figure 3:**
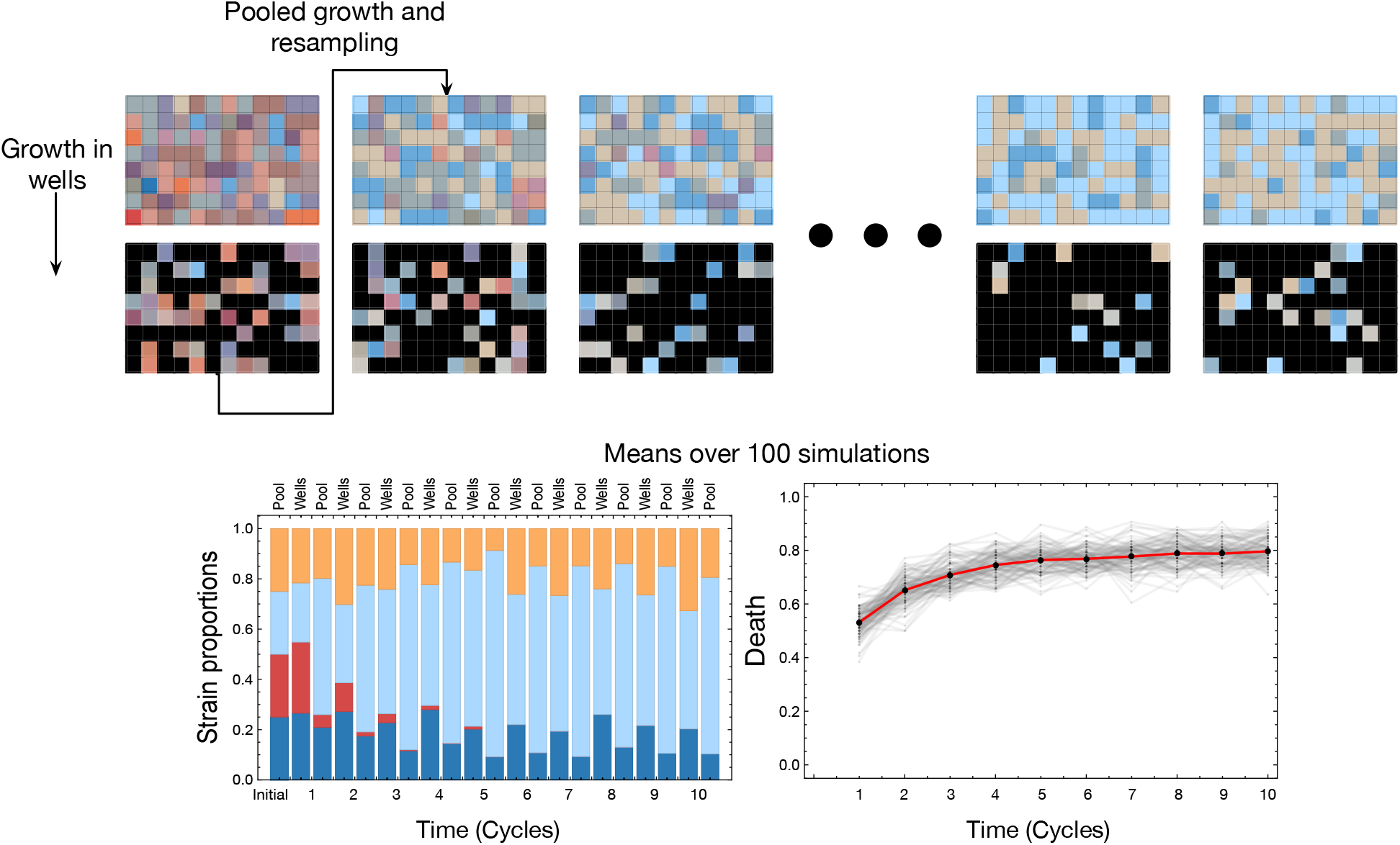
Perfect match. As with tidal cycles in intertidal zones, the strains experience two environments periodically. A pool phase which reflects the growth dynamics as governed by the rates given in Fig. 1 and the wells phase. The wells have a carrying capacity of *K* = 10 and are inoculated with small communities sampled to fill *K/*2. The strains grow with a different growth rates in the wells (as a combination of the community structure and the well ecology) *R*_*i*_(*E, S*) (as calculated in Fig. 1). If a community is not viable then all strains in that community experience the same negative growth rate *d* = −2. For the illustrative example above the time the communities spend in the pool phase is short as compared to the amount of time spent in the wells *t*_*pool*_ = 2, *t*_*wells*_ = 20. As shown in the top panels, the strains cycle between the pool and the well phase. Over time the global composition of the population changes (example of a single run in the barchart and a mean over a 100 runs bottom left) affecting the number of possible ecologies it can survive (given by the mean observed death in the 96 wells, bottom right). All the runs start with all strains present in equal proportions (0.25).

### Ecological variations

The time a community spends in a well is crucial in deciding if the population reaches appreciable densities by the end of the time slot. However, the carrying capacity of the well is a physical constraint that needs to be considered as a deciding factor in estimating the eventual frequencies of the strains transmitted to the pool into the next cycle. In the following sections, we explore the effect of both the time spent in the well-phase of the cycle and constrained carrying capacity.

#### Pool-wells Timescales

The growth rates of the strains are a product of the community composition and the ecology. Thus, as described above, the strains have different growth rates in the pool and the wells. The amount of time the communities stay in the wells versus in pools drives the eventual strain abundances in the whole system. We observe that when the strains spend a sufficiently long time in the pool phase, *t*_*pool*_, the effect of the fitnesses in the wells phase of the cycle is negligible (Fig. 4). Even though the growth rates in the wells are a complex combination of the community composition and the ecology, if the time spent in the wells is sufficiently short, then only the pool growth rates determine the population equilibrium. Essentially the effect of the structured ecology would then be negligible and inconsequential to the long term dynamics of the system. The emergence of complex interactions between the constituent strains is then not observed since the simple single peaked fitness landscape drives the dynamics in Fig. 1.

**Figure 4:**
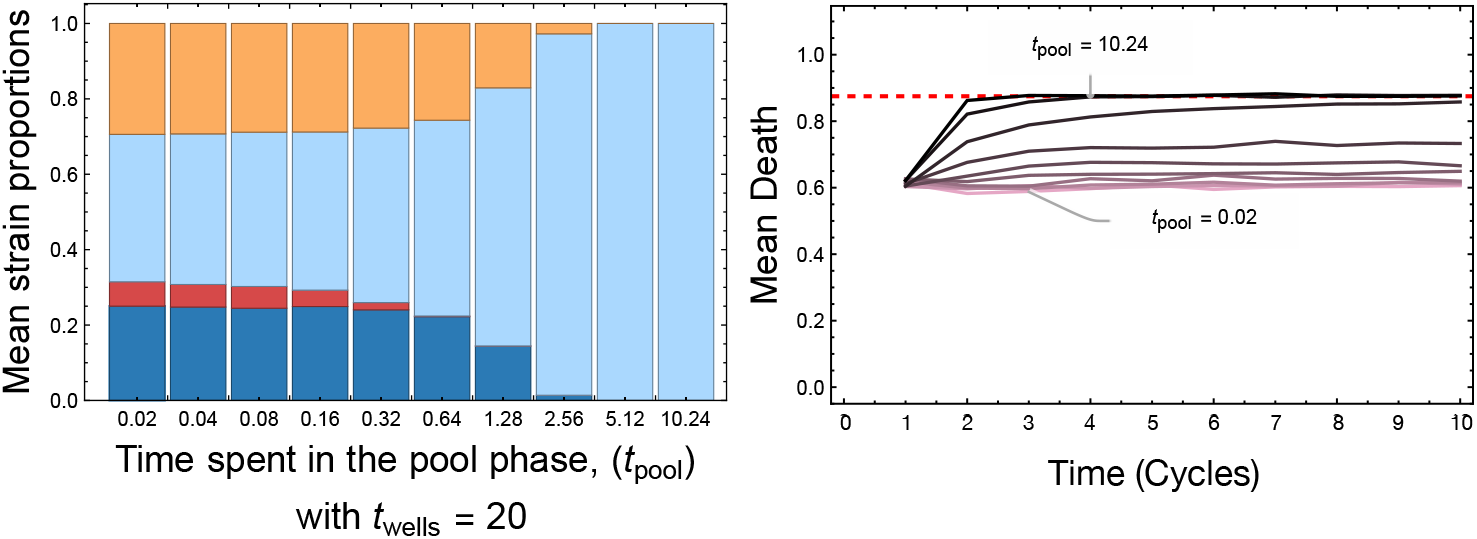
Long term ecological dynamics for different timescales of pool-wells. We assume that the time spent in the wells phase is *t*_*wells*_ = 20. We explore scenarios where the time spent in the pool *t*_*pool*_ is orders of magnitude lower than *t*_*wells*_ or equal. From Fig. 1 we expect that when the time spent in the pool phase is much larger than the one in wells, the *HIS↑* will take over the population. We observe the same already when the time spent in the pool phase is one-fourth (*t*_*pool*_ = 5.12) of *t*_*wells*_. As the time spent in the pool phase reduces with respect to *t*_*wells*_, the equilibrium frequencies reflect the relationship between the growth rates of the strains in the wells in which still the *HIS↑* is dominant but not exclusive (see. Fig. A.1). The mean death (mean over a 100 runs for each *t*_*pool*_) shows that as the population is dominated by *HIS↑*, there is less variability in the communities generated and therefore most of the sampled communities will die in the heterogeneous ecologies. The parameters used for the numerical runs are, *K* = 10, inoculum size = *K/*2, *d* = −2, *t*_*wells*_ = 20. Each of the bars are a mean of a 100 runs where each run begins with a random initial strain proportions.

#### Dependence on well carrying capacity

If the time spent in the well-phase is not a constraint, then final strain proportions are dictated by the carrying capacity of the wells. Differences in strain densities due to varying carrying capacities is exemplified in Fig. 5. Starting from the same initial strain proportions and densities, for a larger carrying capacity Fig. 5 (second column), we observe *HIS↑* taking over the well. Thus when sampling based on pool representation, *HIS↑* will be overrepresented. This overrepresentation of one strain increases the likelihood of extinction in the next generation due to a lack of diversity. Although the lack of produced metabolites can be substituted by viable ecologies, the slight increase in wells that show death captures the effect of the increased carrying capacity. When *HIS↑* takes over not only the well but eventually the pool, then the only environments that can support the strain are {1, 1, 0, 1} and {1, 1, 1, 1}. Since each environment has the same probability of being realised, as previously mentioned, the probability with which a randomly selected well will go extinct is 0.87. Even for a finite sample of 96 wells, we approach this mean death when the environments are equally likely.

**Figure 5:**
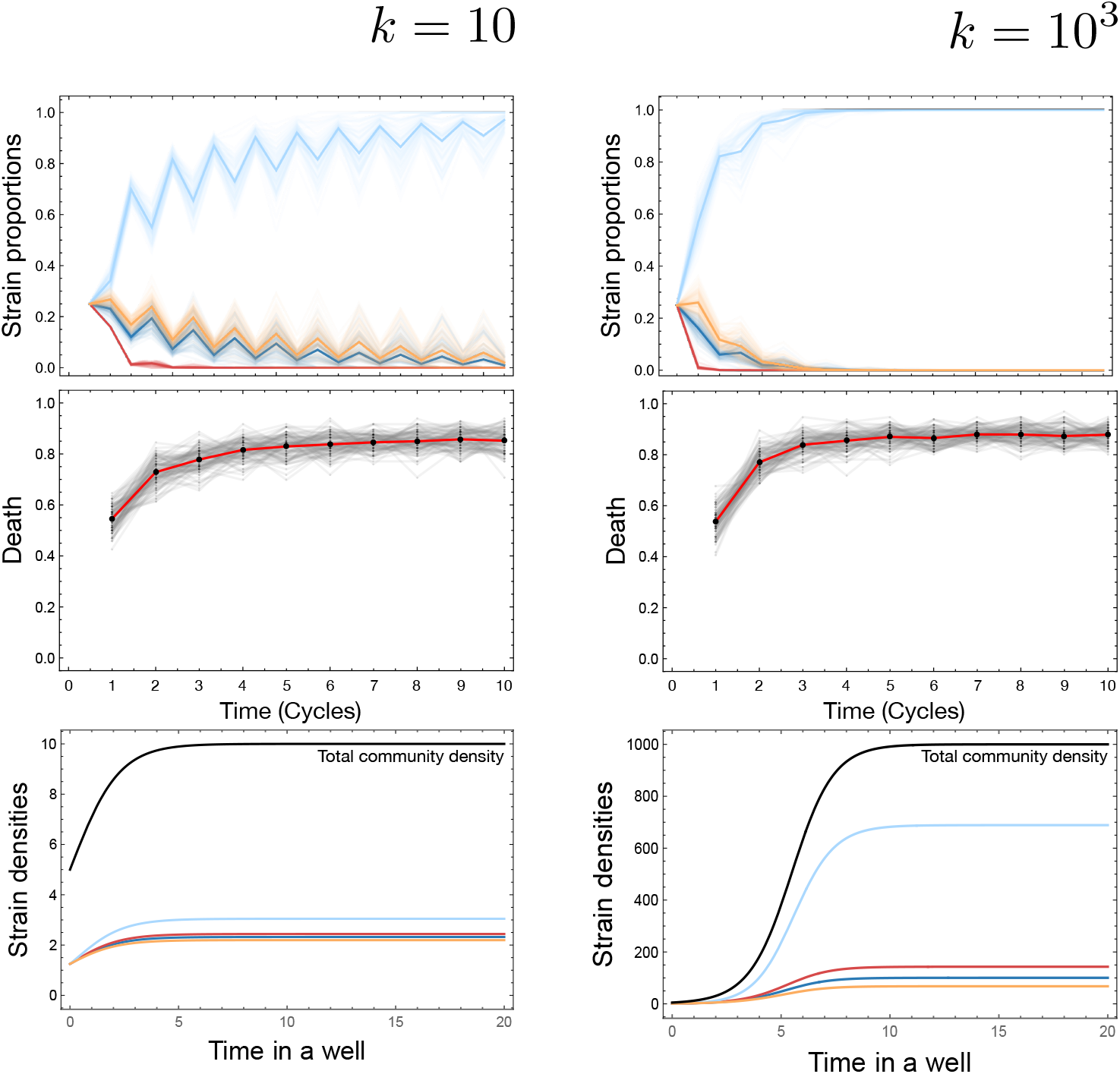
Dependence on carrying capacity of a well. The carrying capacity of the well translates into the population size that a community can achieve during the time spent in the well. The result of two different carrying capacities can be seen in a different in the time required for the *HIS↑* to take over the global population. This also elevates the death observed at the community level as the dominance by one strain reduces resilience. The strain proportions and the mean death are averaged over a 100 runs where for each run all strains are present in equal proportions (0.25). To visualise the precise effect of the carrying capacity we have plotted example dynamics from a particular well (poor ecology with no metabolites) comprising of a community consisting of all strains in equal proportions. The growth rate for this setup extrapolated from the experiments (Fig. A.1) is {0.804, 0.869, 1.157, 0.732} for (*ADE↑, TRP↑, HIS↑, LYS↑*).

However, our assumption that nutrients are available uniformly over the ecological space and hence the assumption of all ecologies being equally probably is highly unrealistic. Furthermore, production of metabolites by each community can also feed into wells and the pool. This ecological feedback of information between cycles leads to our next section, thus completing the loop between the strains and their environment.

### Ecological feedback

#### Niche availability

There are sixteen possible ecological conditions given the four metabolites that can be mixed and matched in our system. For the wells, the ecologies are chosen randomly from these sixteen possibilities in equal proportion. Taking the realistic diversity of nutrients in space and time, we encapsulate this stochasticity in the ecologies by changing the probability distribution used to draw the random ecologies. While earlier all sixteen possibilities were equally likely, now we use a Poisson distribution over the discrete space of the number of ecologies with different means Fig. 6. We drew from three ecological distributions of the 16 possible nutrient combinations: skewed towards single or low numbers of nutrients, skewed towards a medium number of nutrients, and heavily over-representing high numbers of nutrients.

**Figure 6:**
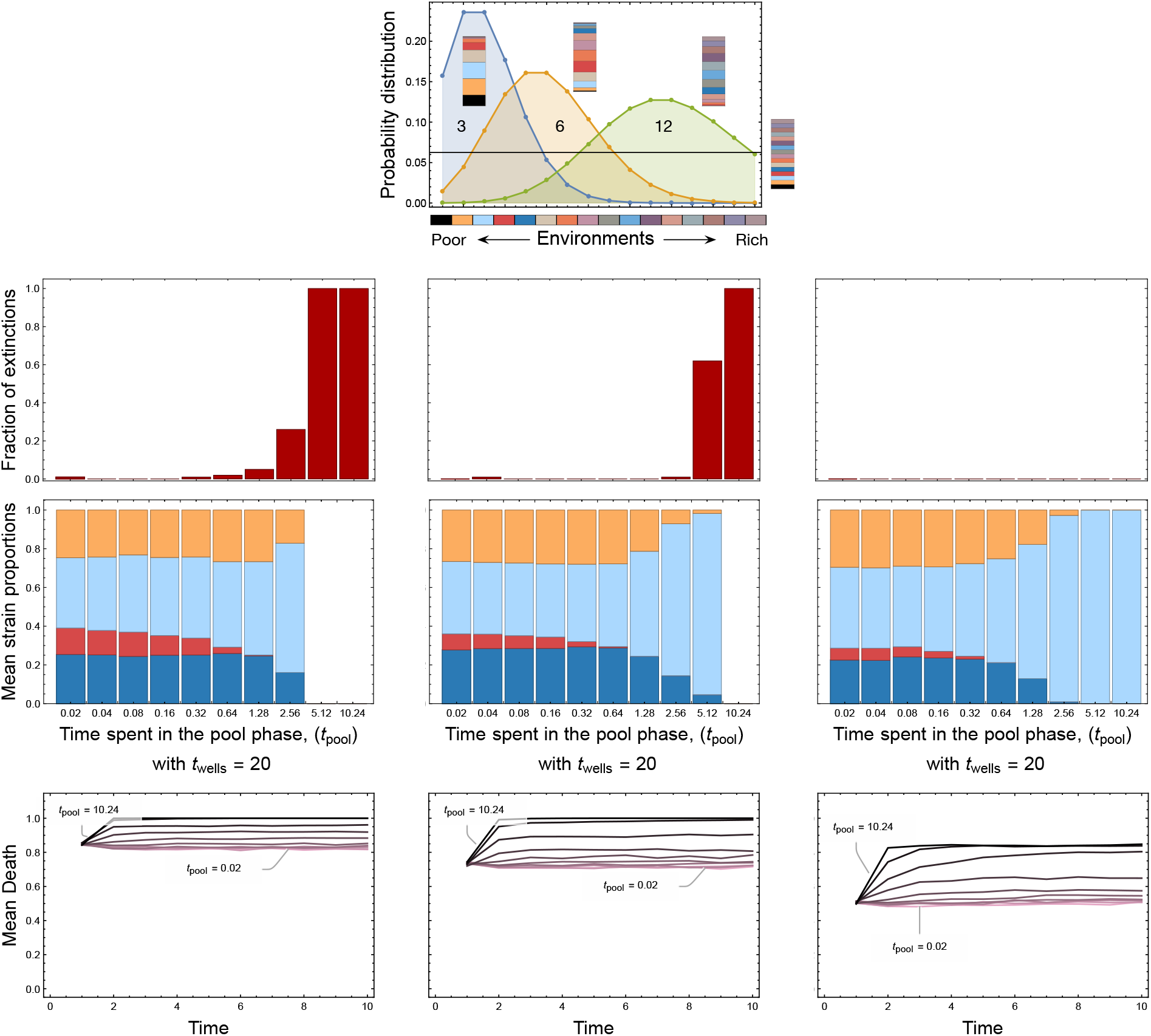
Long term dynamics for different ecological distributions. Until now we had assumed that the sixteen possible ecological variants as seen in Fig. 2 were equally likely. Here we relax this assumption and draw the likelihood of observing a particular ecology with a probability drawn from a Poisson distribution with different means. In the set of panel denoted by A we see the final mean equilibrium frequencies (averaged over a 100 runs) after 10 cycles of pool and well phases. The time spent in the wells is set to *t*_*wells*_ = 20 and the time spent in the pool is varied. The rows are the different environmental conditions where the ecologies follow Poisson distributions with different means as denoted in Panel B. In all cases, the *HIS↑* strain reaches fixation as *t*_*pool*_ increases. We show the mean observed death for the different *t*_*pool*_ which reaches a maximum given by the inverse of the probability of observing the only two permissive environments for *HIS↑* namely {1, 1, 0, 1} and {1, 1, 1, 1}. It is thus possible to estimate the maximum observable death when *HIS↑* has taken over the system for different environmental distributions as given in C. Hence with increasing Poisson mean, as the distribution skews towards the right, we see an over representation of the two conducive environments (denoted by environment 14 and 16) allowing the survival of the system.

#### Niche stabilisation

Organisms typically not only find a niche but are also able to construct it. We have not yet considered the impact of the strains on the ecology itself. Since each strain produces a metabolite, environmental concentrations of metabolites will change. The compositional changes in the strain population thus directly affect the metabolite distributions. The metabolite distributions will ultimately affect the probabilities of observing the different possible ecologies. Hence, the changed metabolite concentrations distort the distribution of ecologies, as seen in the next generation of the wells phase. We take as an example the case where the difference in the timescales between the pool and wells phase is of an order of magnitude (*t*_*wells*_ = 20 while *t*_*pool*_ = 1.28). In this case, the *HIS↑* is overrepresented, with almost 87% of the wells going extinct on an average. We can allow metabolic feedback to begin at different time points in the season. We show in Figure. 7 that if we start the feedback later in the cycling, then the *HIS↑* has already reached appreciable frequencies. Therefore, in the next cycle, most of the ecologies in the wells will have an overabundance of histidine and minuscule amounts of other metabolites. This skew in ecologies hinders the growth of the most abundant strain *HIS↑* itself and leads to higher death in the whole system.

**Figure 7:**
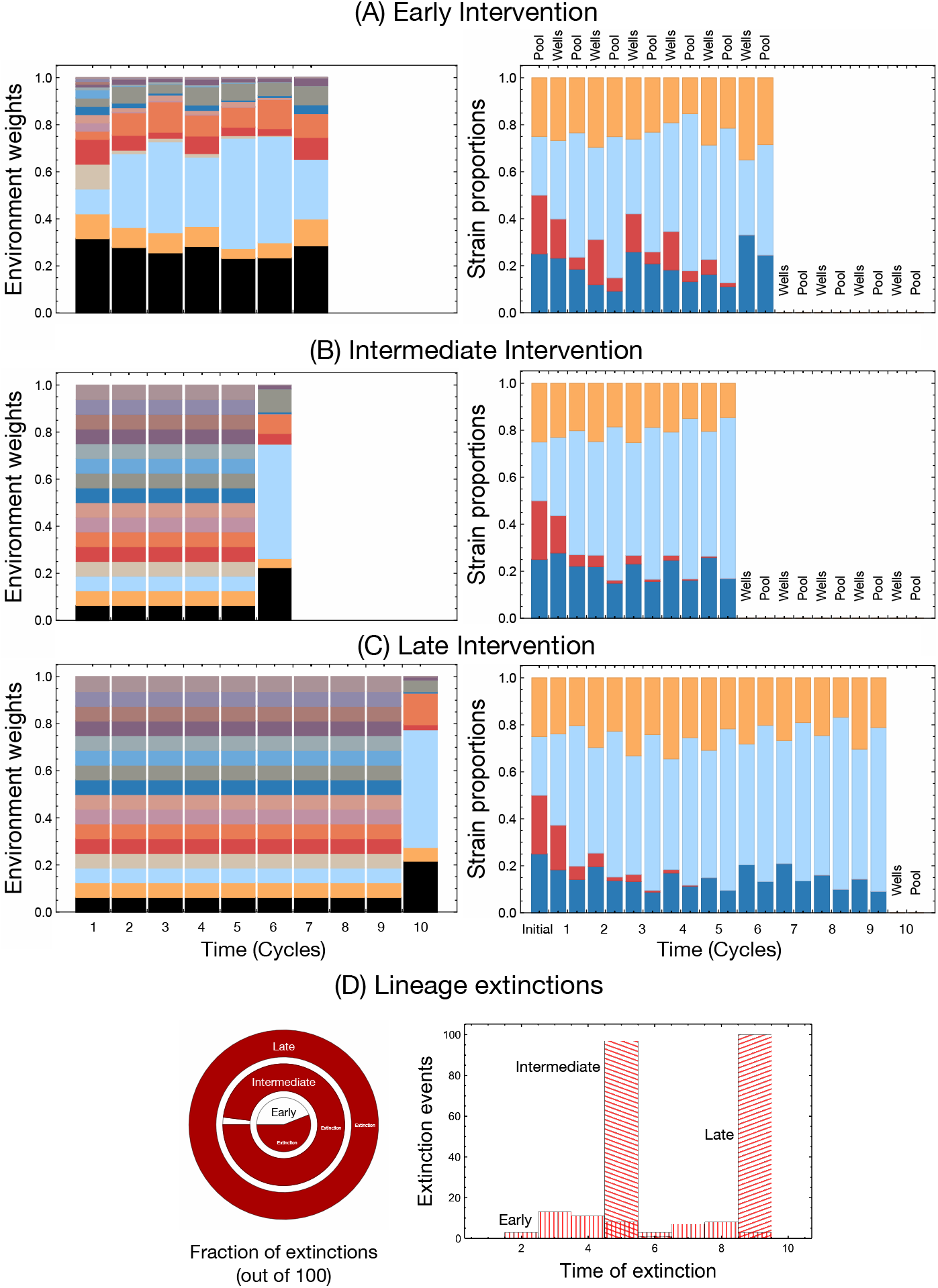
Population dynamics under ecological feedback of metabolites. Up to now we assumed that the ecological cycles exclude the metabolite transport from one cycle to the next. However, focusing only on inter-strain dynamics provides only a partial picture. Here we show the results of allowing for a connection between community population dynamics and and ecology with respect to metabolite concentrations. From Fig. 4 we focus on a particular example where the time spent in the pool phase is set to *t*_*pool*_ = 1.28 and *t*_*wells*_ = 20. The resulting pool frequencies drive the environmental sampling for the next well cycle. We highlight the distributions from which the ecologies chosen (at each well step) on the left and the resulting dynamics of a single run on the right. If we allow for feedback process to start right from the beginning then there is a higher possibility that the lineage survives (A) as opposed to starting the intervention later (C). For the three interventions, the fraction of extinctions as calculated over multiple (56,98 and 100 out of 100) runs is denoted in the pie chart at the bottom with the time distribution of extinctions in the right plot.

#### How to avoid collapse?

Methods of avoiding collapse can also come from ecological and evolutionary processes. *Ecologically*, if feedback is already possible from the beginning of the ecological cycling, then the overgrowth of one strain is curbed (Fig. 8 top row). The whole system then has a higher probability of surviving (lower fraction of extinction), and the slight risk spreading over time. Alternatively, *evolutionarily*, we show that as growth rates of the strains equalise then, we also see a decrease in the number of extinctions (Fig. 8 bottom row). For both the ecological and evolutionary approaches of avoiding extinctions, almost all the lineages survive early interventions, and the interventions at intermediate or later stages are now less lethal. Designing an ecological structure where the pool time is reduced or engineering strains with only slightly different growth rates could enhance the survivability of artificially designed communities and keep them stable against environmental fluctuations and feedback processes.

**Figure 8:**
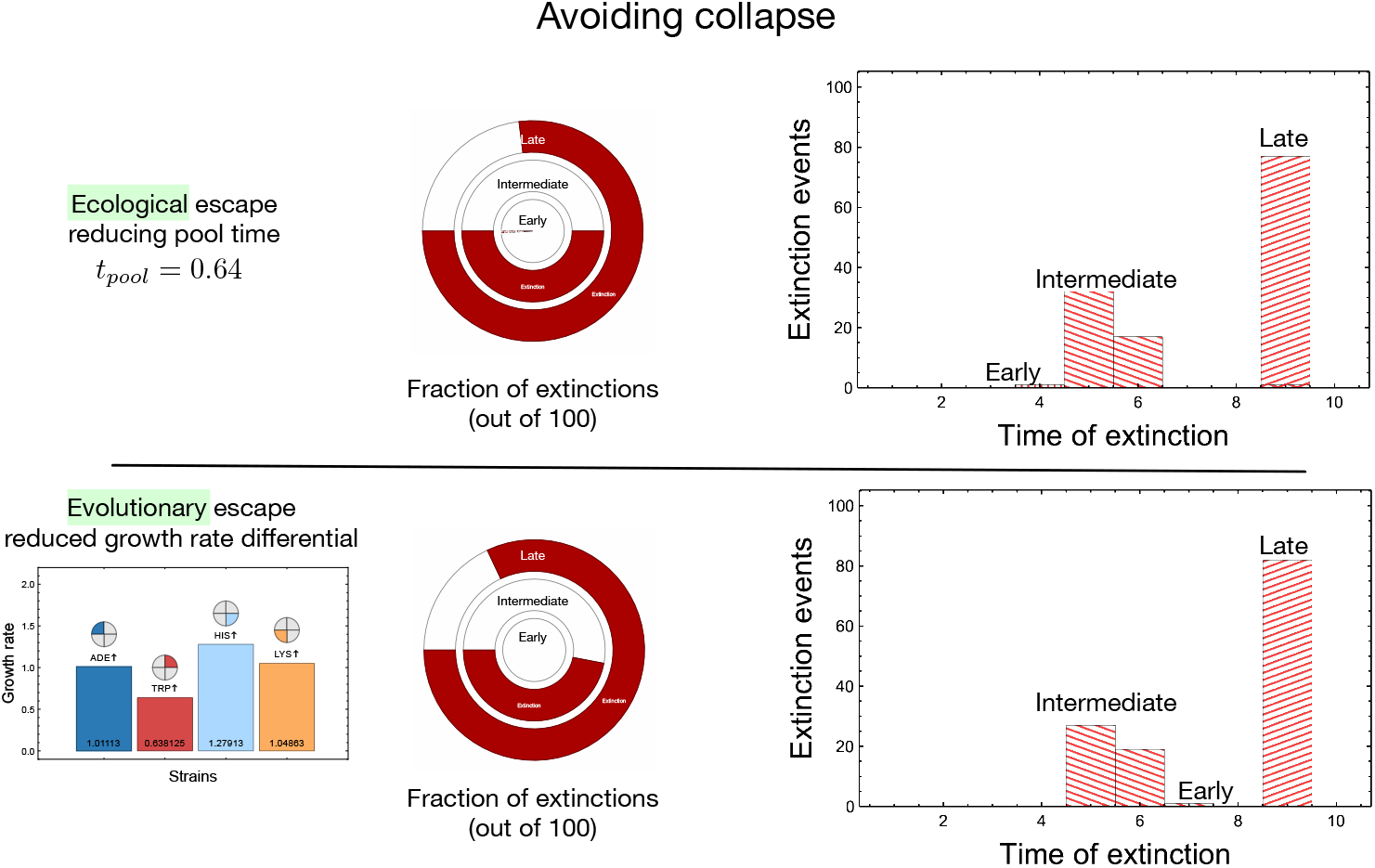
Avoiding lineage collapse under ecological fluctuations. Both ecological and evolutionary processes could allow us to avoid the collapse of communities at the lineage level. From Fig. 7D we observe that for intermediate and late interventions, all lineages go extinct. In the top row we show that if the amount of time spent in the pool phase is shorter (half as that of Fig. 7D) then we can ecologically rescue a substantial fraction of the lineages in both the intermediate and late intervention treatments. In the bottom row, we show that rescue is possible if the growth rate of the strains are quantitatively less different from each other. For example we now bring the the growth rates closer to the mean of all strains by 50% as compared to Fig. 1. This change maintains the differences between the growth rates qualitatively but already we see a decrease in the number of extinctions. For synthetic systems, we envision that both of these processes can be engineered.

#### Evolutionary fate

We have previously discussed a two strain synthetic system [28] based on the works of [22]. The engineered strains lacked the production of adenine or lysine. A code could define the strains: {*m*_1_, *m*_2_} where the production of a metabolite (*m*) can be represented by on 1 or off 0. Thus the wild-type is denoted as {1, 1} and the two constructed strains are {1, 0} and {0, 1}. The latter two can grow only in a rich environment where the unsecreted metabolite is present or in the presence of each other where they support each other via the engineered mutualism. However, binary systems are typically exceptional cases of complex systems [32]. We augmented the two-strain systems with different interdependencies, now tryptophan and histidine coming from [23]. Engineering different levels of interdependencies with four metabolites can be represented by {*m*_1_, *m*_2_, *m*_3_, *m*_4_} where again we have *m*_*i*_ = {0, 1}. Thus 16 different strains are possible. Our study focused on the extreme four, where the strain can produce only one metabolite.

The costs and benefits of acquiring and losing essential genes can be highly complicated and environment-dependent [33, 34]. According to the Black Queen hypothesis, the loss of essential genes for leaky products can be feasible if the environment can compensate. Furthermore, gaining new genes might not always come at a growth cost. A thorough analysis of the cost-benefit structure in the light of the ecology is, therefore, necessary for systems that can incorporate the evolution of these interdependence traits [20, 35]. The possibility of tuning the strengths of interspecific interactions can, in turn, also affect the observed biodiversity levels in evolving communities [36].

## Discussion & Conclusion

Cyclical variation in ecology affects interactions, population dynamics, abundance and richness in a range of systems [37, 38]. Despite this knowledge, the fundamental role of environmental periodicity is understudied [39]. We conceived a stable evolutionary system with fixed functional diversity operating on short timescales to understand how cyclic ecological processes influence abundance and community formation of organisms in a biological system. This system showed how cyclical fluctuations in resource availability and population-resource feedback loops shape network interactions. Our model embeds a four-strain interaction network in an environment cycling between structured and unstructured phases to explore how ecology shapes community formation, cohesion and persistence. We established how cyclic nutrient availability generates spatial and temporal heterogeneity capable of altering resource exploration, modifying the selection on ecologically relevant interactions (from cooperation to competition) and potentially serving as the basis for evolutionary dynamics.

Our system comprises cyclical compartmentalisation of replicators and resources. Essential resources, such as nutrients, water and space, often fluctuate in regular cycles linked to major forces like the movement of the earth. Hence, living organisms are subject to changes in conditions that will affect their survival and reproductive potential. The process is, however, more general, extending to prelife [40]. At the level of prebiotic molecules, the structure of alkaline hydrothermal vents is pockmarked with the essential nutrients of autocatalytic reactions flowing in and out through them in a heterogeneous manner [41]. Alternatively, for the RNA world hypothesis, transient compartmentalisation and dry-wet cycles of the warm little ponds are postulated to be necessary for polymerisation to generate complexity [40, 42]. At the level of individual organisms, species adapted to live on rocky shores are subject to periodic changes in conditions caused by rescinding tides, which can lead to increased mortality and modify population structure and species abundance [43, 44]. Although cyclical events usually occur alongside a continuum, compartment-based models can provide a better alternative to its study. We used resource availability and defined carrying capacity to create distinct environmental compartments. Each cycle (compartmentalised and homogeneous) acted as an environmental sieve that led to changes in population density and frequency, which would be carried out to the next cycle (Figure 3).

Our four-component experimental system exhibited substantial growth differences between engineered strains. Therefore, long growth periods in the nutrient-rich pool phase, without interdependencies, exacerbated these growth differences and reduced the representation of weaker strains. Thus, compartmentalisation that limits such unrestricted growth reduces the impact of strain growth differences (Figure 4). However, increased well-size can offset this dependence on compartmentalisation (Figure 5). Unremarkably *HIS↑* dominance increased with the carrying capacity of the wells as this provided greater opportunity to leverage growth differences. Our implementation of cyclical compartmentalisation, the pool-well-like dynamics, are highly prevalent in naturally occurring systems subject to regular occurrences such as tides, seasons or monsoons. In the compartmentalisation of transient estuaries and intertidal zones, niche size and timing are widely known to alter ecological and evolutionary dynamics. For example, trait evolution is linked to compartment size in transient estuaries [45]. Furthermore, temporal compartmentalisation has led to different ecotypes of the marine midge Clunio [46]. However, in these real-world situations, niches within each compartment are not random. They are shaped by existing factors and thus are a subset of the broader systems. These dynamics drive niche size and nutrient presence and thereby alter the strain abundances and the eventual ecosystem stability [47, 48].

Fixed interactions and nutrient limitations highlighted that the successful establishment of a given niche is highly dependent on pool composition in previous cycles. We also showed that seasonal niches could mediate functional diversity and influence the ecological dynamics between the interactive communities [49, 50]. Suppose the distribution of the metabolites is not equal in the pool. In that case, the sampling of the environments in compartments will also not be uniform. Allowing richer environments to be more likely, we unsurprisingly see fewer lineages going extinct. However, the persistence of lineages does decrease the diversity (Figure 6 right column). For instance, an organism’s functional role is maximised with limited resources and the interdependence between cooperating strains increases (Figure 6). High resource availability had the opposite effect. It eroded interdependence and promoted competition between strains [51, 52], which led to a population expansion, followed by dominance, reducing viability in the well phase, which led to an eventual collapse of the lineage. This feedback quickly resulted in strain extinction and an increased death rate (Figure 6). Disruption of these normal cyclical cycles in natural environments can result in ecosystem collapse. For example, dead zones are areas of water bodies where aquatic life cannot survive due to low oxygen levels. One of the most common causes of dead zones is eutrophication, where excess phosphorus and nitrogen from fertiliser run-off or sewage cause an excessive algal bloom, which depletes other dissolved nutrients in the water and oxygen, leading to the death of other aquatic organisms and eventually, the algae [53–55].

Varying the availability of nutrients in our pools resulted in a trade-off between maintaining diverse communities and overall survivorship (Figure 6). In niches with lower diversity, greater inter-strain dependency existed. Rapid extinction was observed with minimal scope for nutrients in any given niche to compensate for the loss of even single strain. Due to our system’s fixed and non-overlapping biological function, each strain is critical to optimal community survival. That effectively makes the weakest strain, *TRP↑*, a keystone species (see what happens if we remove all TRP containing environments in Fig. A.3). Essentially all aspects of a given biological system are impacted by a continuum of biological processes occurring at every level of the organisation. Engineered interdependencies enforces this relationship idea and allows us to dissect biological concepts like keystone species. It was impossible to predict the ecological impact of reintroducing wolves to yellow-stone park due to the enormous number of uncharacterised interactions and the subsequent flow-on from these interactions [56]. However, with defined fixed interactions, we are in a position to model ecological outcomes in simplified systems. For example, the explicit linking of the loss of weaker but critical species to ecological collapse.

As discussed, factors that help preserve strain diversity result in an overall lower death rate and stave ecosystem collapse. However, we sought to explore further the evolutionary and ecological factors that prevented collapse. We allowed each pool-well cycle to influence the nutrient composition of the next cycle based on strain composition (Figure 7). In this case, strain-induced environmental changes become a model for the Allee effect [57, 58]. An early intervention, allowing inter-cycle feedback from the first cycle, maintained all four strains in most instances. The low frequency of any given strain reduced the corresponding nutrient availability and lowered overall population fitness [59] but provided an advantage to the underrepresented strain. In previous interactions of our model, complete extinction, even if only a single strain survived, was impossible due to the equal provision of nutrients. However, with intermediate and late inter-cycle interventions, complete extinction was immediate. Confirming the keystone species dynamics, the loss of any single strain always results in extinction with the inter-cycle interactions.

Ecological and evolutionary dynamics are possible means of avoiding ecosystem-wide collapse (Figure 8) [60–62]. Ecologically if the amount of time spent in the pool phase is reduced, the strain with the highest growth rate will not have enough time to outcompete the others. Evolutionarily the lineage could be rescued if the growth rate of the individual strains becomes comparable. Allowing for variability in growth rates may be the first step toward explicit evolutionary processes. Our simple experimental and theoretical model allows us to clarify and understand the effects of these two processes separately. This simplicity allows us to probe scenarios from prebiotic chemistry at the inception of evolution to modern living systems where both of these processes act in tandem.

Our model allowed dissection of ecological impact via static biological functions of strains resulting in predictable interactions within a community. Experimental and theoretical systems that permit the biological function to evolve can provide enormous insight into the dynamics that shape communities [20]. It can, however, be challenging to untangle causality in genetic and environmental factors [63]. Thus fixed biological functions are often used to re-move the role genetics play [17, 22, 23]. Incipient evolutionary processes need to necessarily exclude the presence of interactions between replicators – from cells to species. One of the methods to impose selection is then via ecological scaffolding, where the ecology selects on a specific property of the collective [64]. In our study, we exemplify the role of ecology but do not assume any property of the community to select for but rather propagate what persists [26]. Persistence will then provide a fertile testbed for evolutionary processes to develop. Endogenisation of cyclical ecological processes such as the development of lifecycles, circadian, and circalunar cycles may provide an evolutionary advantage to communities.

Ecological networks can be highly variable concerning their constituents [65–67]. However, our current model has only a limited capacity for modularisation. Future work facilitating a greater understanding of complex eco-evolutionary network dynamics requires greater modularisation by adding additional nodes and subnetworks. A logical first step is to develop experimental and theoretical systems that reflect greater strain diversity. Specifically strains that represent all 16 combinations of function, from sharing none of the metabolites to sharing all four. This system allows the formation of multiple possible survivable consortia. Additionally, this developed system will help test if greater diversity will allow for greater stability of lineages. We will thus be able to advance our understanding of nested ecological dynamics, consortia expansion and cross-scale interaction.

This theoretical framework is a part of a larger ongoing project on the evolution and ecology of synthetic communities [22, 23, 28]. An application of an extended consortia system, and a key driver behind the development of this model, would be in predicting the outcomes of ecosystem interventions. For example, synthetic microbial communities with complex biological functions are being developed for fields in translational biology such as conservation, health and exploration from bioremediation, and biodiversity restoration to space mining [68–71]. We have previously defined consortia with set inputs and outputs as swappable “ecoblocks” in synthetic microbiology [27]. However, the role of ecology where such ecoblocks might be released is vital but so far neglected. Assessing the viability of such consortia using dynamic ecological principles as here will help design efficient synthetic consortia or via in-silico driven artificial selection [73]. Once the communities have achieved their goal, self-limitation through community extinction might be a desired feature and not a bug [27].

To say nothing in biology makes sense except in the light of evolution perhaps does a disservice to biology. It might be fairer to present this idea in a more expanded fashion - nothing in biology makes sense except in the light of evolution illuminated by ecology. We have highlighted the power of cycling ecological processes to form and select communities for persistence, focusing only on the growth parameters [74]. Given the ubiquity of cycling processes, as described earlier, the implications of our work range across the hierarchies of systems in prelife and life, while the information can be used to design specific applications in translational biology.

## Acknowledgements

This work was supported by JSPS KAKENHI Grant Number 19K06795 to JAD and CSG. CSG acknowledges support from the Max Planck Society. MV & JAD were supported supported by the Okinawan Institute of Science & Technology and Monash University. We would like to thank the laboratories of Dr D. Greig and Professors W. Shou and A. Murray for providing yeast strains.

## Author Contributions

CSG developed the theoretical model, performed the experiments, analysed the results and wrote the paper. MV performed the experiments and wrote the paper. JAD developed the yeast system, performed the experiments, analysed the results and wrote the paper.

### Data Availability and Analysis

All code and preliminary plots are available on GitHub at https://github.com/tecoevo/ecoblocs.

### Code availability

https://github.com/tecoevo/ecoblocs.

## A Methods and Materials

### Experimental Materials

#### Saccharomyces cerevisiae Strains

The synthetic system is composed of four metabolite- overproducing strains developed in the w303 strain background https://github.com/tecoevo/ecoblocs/blob/main/strain_creation/strain_list.csv. Oligo primers used in this study are listed in https://github.com/tecoevo/ecoblocs/blob/main/strain_creation/primer_list.csv. Strain development relied on creation of a progenitor strain, from w303-1, with auxotrophies for adenine, lysine, tryptophan and histidine and the insertion of feed back resistant (FBR) mutations that result in the over production of either lysine, tryptophan or histidine. Specifically, the KanMX, derived from plasmid p0003 (Addgene 44901), marker was inserted into the *HMLα* loci using primers P0114 and P0115 to prevent mating type switching. *ADE4* was disrupted with *CaURA3* amplified from plasmid p0011 using primers P0118 and P0119. *LYS2+* was disrupted using function copy of *LEU2+* amplified from W303-2 genomic DNA using primers P0123 and P0124. This resulted in the Progenitor strain.

The *ADE↑* strain was created by replacing *ade4::CaURA5* with *ADE4(PUR6)*, the FBR version of *ADE4*, amplified from yeast strain WS950 using primers P0155 and P0156. *URA3+* amplified from WS950 using primers P0069 and P0070. The *HIS↑* strain was created by insertion of (*HIS1FBR, HIS3*) from plasmid p0050 into the *HIS1* locus using primers P0151 and P0152. The *TRP↑* strain was created by insertion of (*TRP2FBR, TRP1*), from plasmid p0044, using primers P0309 and P0310. The *LYS↑* strain was created by insertion of (*LYS21FBR, LYS2*), from plasmid p0045, using primers P0312 and P0313.

#### Plasmids

A list of plasmids used in this study is provided in https://github.com/tecoevo/ecoblocs/blob/main/strain_creation/plasmid_list.csv. Gibson assembly was used for plasmid creation (NEB E2611L). Plasmids p0003 and p0011 were ordered from AddGene (plasmid IDs 44901 and 44650 respectively). The plasmid pUC19 (NEB N3041S) was used as a backbone for all plasmids constructed. Plasmid p0050 (*HIS↑*) was created by cloning *HIS1* and *HIS3* using primers P0145-P0148 into pUC19. *HIS1* was mutated to the feedback resistant form described in [30] using primers P0149 and P0150. Plasmid p0044 (*TRP↑*) was created by cloning *TRP2FBR* and *TRP1* using primers P0157-P0160. *TRP2FBR* was amplified from yeast strain yMM65 [23]. Plasmid p0045 (*LYS↑*) was created by cloning *LYS21FBR* and *LYS2* using primers P0161-P0164 into pUC19. *LYS21FBR* was amplified from yeast strain WS954 using primers P0161 and P0162.

#### Media

Synthetic complete (SC) media was made from dextrose, FORMEDIUM yeast nitrogen base and an appropriate synthetic complete dropout supplement (Kaiser). Amino acid free minimal media (SC-aa) was made as above without synthetic complete dropout supplement. Yeast-extract, peptone, dextrose (YPD) media was made using chemicals from BD Diagnostics or Sigma.

## Experimental Procedures

### Culturing

All cultures were grown at 30 °C in an orbital shaker at 200RPM. Each experiment was performed in triplicate starting from a single individual colony. Experimental cultures were generated by selecting individual colonies from a streak plate and growing each colony in 5 mL SC medium for 48 hours. These cultures were pelleted, washed twice with SC-aa and resuspended in 5 mL of SC-aa. They were then grown for an additional 24 hours. This allowed the culture to reach a carrying capacity of approximately 5 *×* 10^7^. These cultures were pelleted, washed and resuspended as above and then were diluted 1 in 20 with SC-aa. Thus, all experiments were at approximately one twentieth of the carrying capacity. Experimental cultures were generated by mixing individual strain dilutions in equal proportions to a final volume of 5 mL. Nutrients were added as needed to a final concentration of 100 µg/mL at timepoint 0. The cultures were sampled to determine growth and strain ratios every 24 hours (including timepoint 0). All experiments were performed in triplicate.

### Cell counts, Strain Ratios & Culture Growth

All experimental cultures were sampled by removing 20µl every 24 hours. Cell counts per millilitre of viable cells were obtained from experimental cultures using the Muse cell counter (Merck Millipore) with the Cell Count & Viability kit per the manufacturers instructions. To obtain strain ratios, samples were serially diluted, played on YPD solid media, allowed to grow for two days and colonies were counted. These plates were then replicated onto solid media that permitted growth of only one strain, SC without lysine, adenine, tryptophan or histidine, counts of each strain were obtained, and ratio calculated. Growth rates were calculated for all pair-wise strain combinations as well as all four strains together. Growth rate was determined as the average growth across the first three days of growth.

### Estimating growth parameters

Growth constants underpin theoretical modeling but are highly complex in biological systems. Although increasingly complex derivatives can be employed to capture the temporal life history of a particular organism, these are approximations and increase the complexity of subsequent models. Moreover, it is impossible to generalize growth rates in response to biotic factors, such as other strains, and abiotic factors, such as nutrient availability, due to the enormous numbers of inter-dependencies that further complicate these complex temporal life histories. As such, while grounding our model in laboratory based experiments, we have taken a high-level approach to the generation of a growth values that summarize these complex dynamics. Laboratory-based growth analysis was conducted for a large subset of growth conditions with missing data estimated from these. Unfortunately, as we no longer have laboratory access, additional measurements have been impossible to obtain.

We determined growth rates of each strain in our four-strain system when either grown together, four strains per culture, and in all pairwise combinations, two strains per culture. To reduce temporal-induced variation, growth values are the average of three days growth. Growth rate was calculated using the formula “(cells at time *t* + 1 - cells at time *t*) / cells at time *t*” for each replicate or each culture. For determining pool-growth we used the 4-strain cultures with growth averaged over three days. This was *ADE↑* :1.028 / *TRP↑* :0.282 / *HIS↑* :1.564 and *LYS↑* :1.103. Triple-wise and varying ecology strain growth rates were estimated by averaging growth rates of each relevant pair-wise and four-strain growth rates. For example, to calculate the *ADE↑* strain growth rate when grown with *HIS↑* and *TRP↑*, the growth rate is the average of the four-strain growth rate and the growth rates from the *ADE↑* / *HIS↑* and *ADE↑* / *TRP↑* pair-wise growth rates. These growth rates are shown in fig. A.1. In addition raw data and growth rates are provided on (GitHub).

### Alternative estimation of growth parameters

Estimating the growth rates of individual strains given the community and the ecology is non-trivial. Due to the lack of experimental data we hypothesize the growth rate from our estimates from a subset of experiments. Since the extrapolation can be done in several ways here we discuss another method and display the results accordingly. We identified no key differences between these methods in terms of the conclusions drawn from this work. This suggested that regardless of how this interpolation was achieved, the influence was negligible as it was still tightly bound by the experimental data. The alternative method, more subjective method, employed a maximum growth strategy. Herein the highest growth rates of each strain were favored in media with a higher degree of supplementation. Where all required nutrients were provided this highest rate was used. In conditions requiring inter-strain cross feeding was required, growth rates from pairwise interactions were employed. However, this did serve to increase overall differences between the strongest strain, *HIS↑*, and the weakest strain, *TRP↑*. The data is available at https://github.com/tecoevo/ecoblocs/tree/main/growth_data

### Keystone

If the environment cannot provide what is essential for survival then the organism that can becomes a keystone of a community that thrives there. We show this intuition by removing all environments that involve tryptophan. *TRP↑* is the slowest growing strain but now it becomes important as it is the only source of TRP for the rest of the strains. As seen in Figure A.3, *TRP↑* can grow to appreciable frequencies and even replace the next slower growing strain *ADE↑*.

**Figure A.1:**
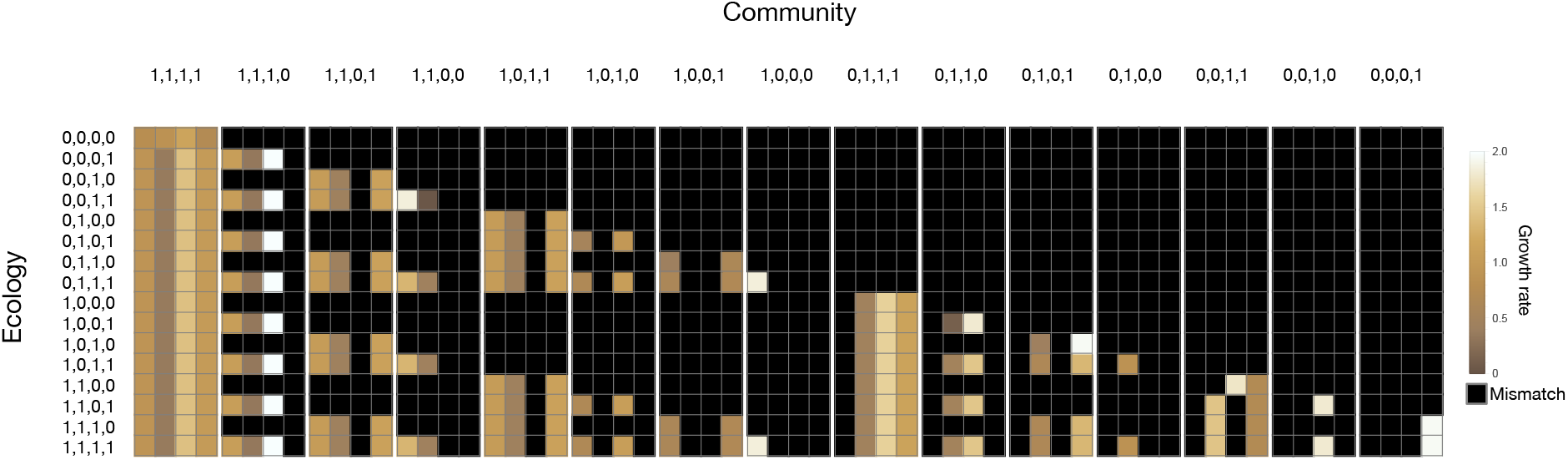
Average strain growth rates. Three day average of pairwise and four-strain growth rates were used to estimate missing community growth rates. This was achieved through the averaging of available experimental data.

**Figure A.2:**
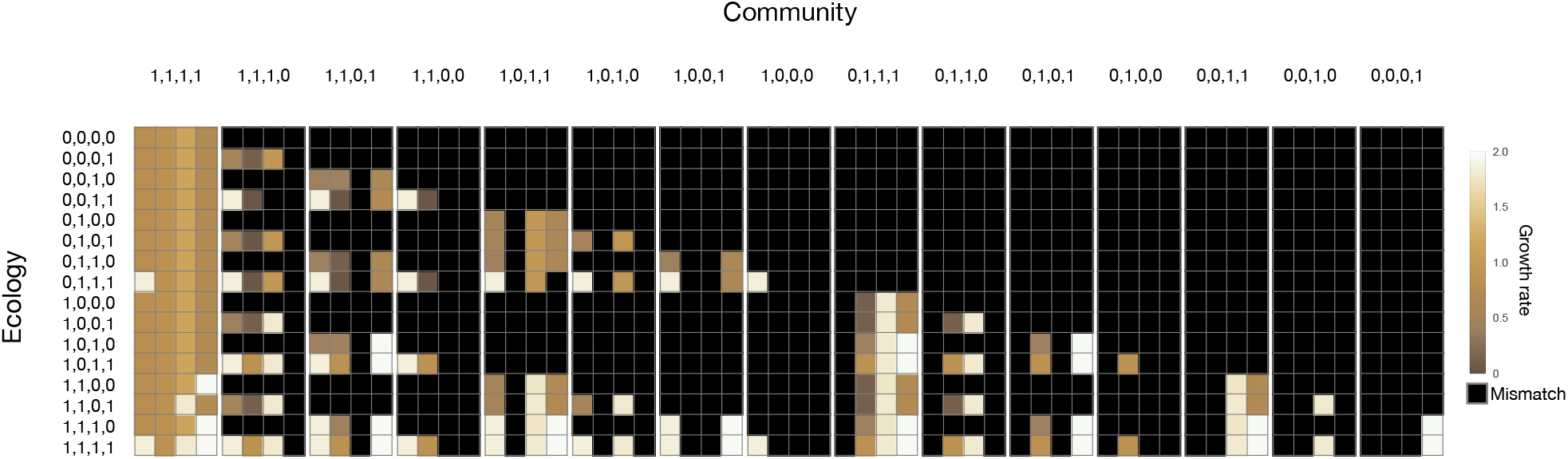
Alternative Strain growth rates. Three day average of pairwise and four-strain growth rates were used to estimate missing community growth rates. This was achieved through selection of maximum strain growth, the highest level achieved, and using the value in condition where all required nutrients were provided. Where inter-strain feeding was required for survival, growth vales from pairwise interactions were employed.

**Figure A.3:**
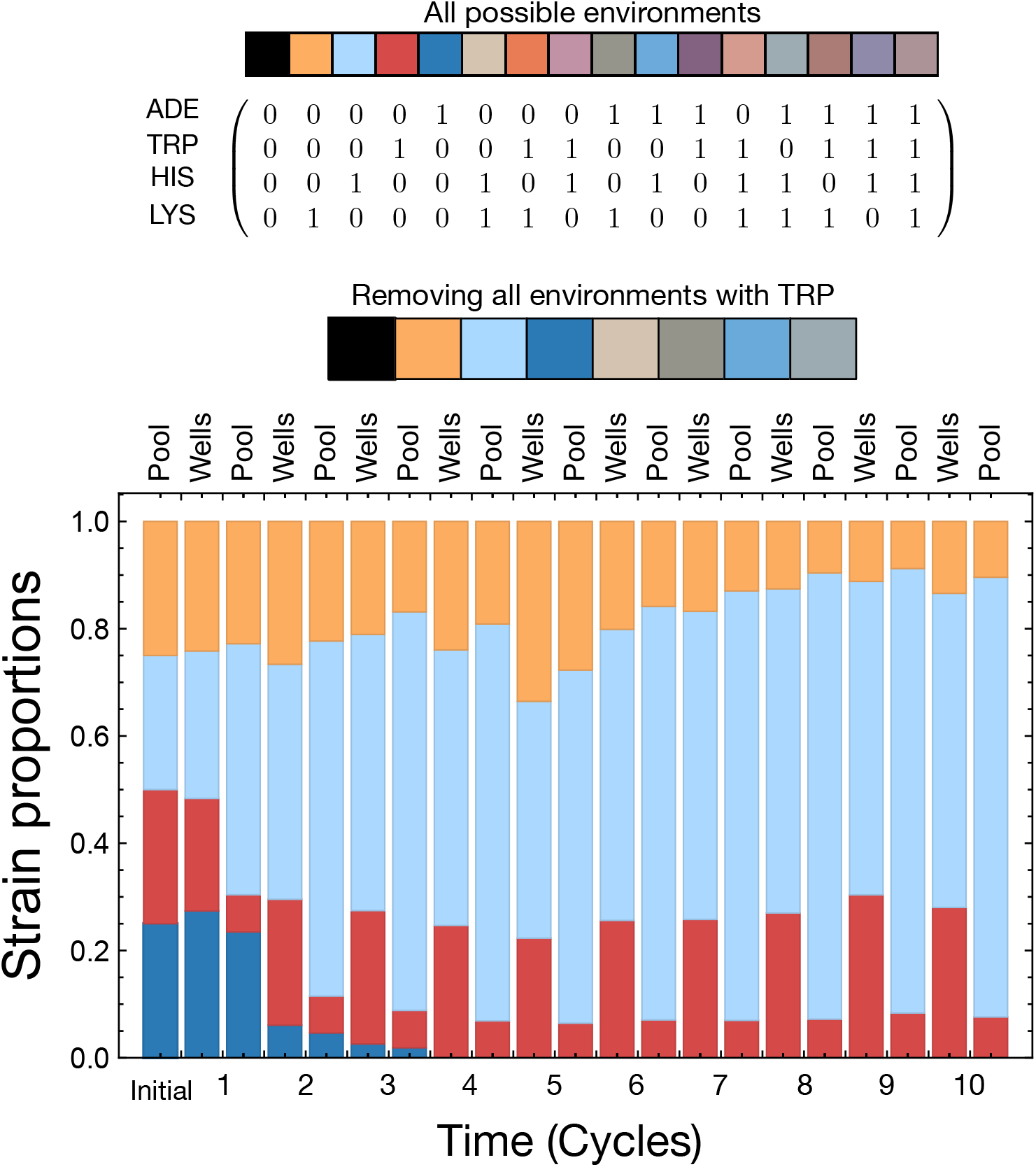
*TRP↑* as keystone. For the same parameters as in Fig. 3, instead of all possible environments being possible if we remove all environments involving tryptophan then *TRP↑* becomes an essential component of the community. *TRP↑* has the lowest growth rate (Fig. 1) and typically goes extinct when all metabolites are externally provided. However, in the reduced set of possible environments (as here), *TRP↑* is invaluable and can even displace *ADE↑* from the system, the next slowest growing strain.

## References

1. Swemmer, A. M. et al. The ecology of drought – a workshop report. South African Journal of Science 114 (2018).

2. Fensham, R. J., Laffineur, B. & Allen, C. D. To what extent is drought-induced tree mortality a natural phenomenon? Global Ecology and Biogeography 28, 365–373 (2019).

3. Ondier, J. O., Okach, D. O., Onyango, J. C. & Otieno, D. O. Interactive influence of rainfall manipulation and livestock grazing on species diversity of the herbaceous layer community in a humid savannah in Kenya. Plant Diversity 41, 198–205 (2019).

4. Xu, X., Medvigy, D. & Rodriguez-Iturbe, I. Relation between rainfall intensity and savanna tree abundance explained by water use strategies. Proceedings of the National Academy of Sciences 112, 12992–12996 (2015).

5. Janzen, D. H. Encyclopedia of Biodiversity, chap. Latent Extinction—The Living Dead, 590–598 (2001), second edition edn.

6. Valiente-Banuet, A. et al. Beyond species loss: the extinction of ecological interactions in a changing world. Functional Ecology 29, 299–307 (2015).

7. Leclère, D. et al. Bending the curve of terrestrial biodiversity needs an integrated strategy. Nature 585, 551–556 (2020).

8. Koh, L. P. et al. Species Coextinctions and the Biodiversity Crisis. Science 305, 1632–1634 (2004).

9. Palmer, T. M. et al. Breakdown of an ant-plant mutualism follows the loss of large herbivores from an African savanna. Science 319, 192–195 (2008).

10. Terborgh, J. et al. Tree recruitment in an empty forest. Ecology 89, 1757–1768 (2008).

11. Anderson, S. H., Kelly, D., Ladley, J. J., Molloy, S. & Terry, J. Cascading Effects of Bird Functional Extinction Reduce Pollination and Plant Density. Science 331, 1068–1071 (2011).

12. Veen, F. J. F. v., Holland, P.D.v. & Godfray, H. C. J. Stable coexistence in insect communities due to density - and trait mediated indirect effects. Ecology 86, 3182–3189 (2005).

13. Martorell, C. & Freckleton, R. P. Testing the roles of competition, facilitation and stochasticity on community structure in a species-rich assemblage. Journal of Ecology 102, 74–85 (2014).

14. McIntire, E. J. B. & Fajardo, A. Facilitation as a ubiquitous driver of biodiversity. New Phytologist 201, 403–416 (2014).

15. Miele, V., Guill, C., Ramos-Jiliberto, R. & Kéfi, S. Non-trophic interactions strengthen the diversity—functioning relationship in an ecological bioenergetic network model. PLoS Computational Biology 15, e1007269 (2019).

16. Cadotte, M. W., Carscadden, K. & Mirotchnick, N. Beyond species: functional diversity and the maintenance of ecological processes and services. Journal of Applied Ecology 48, 1079–1087 (2011).

17. Momeni, B., Chen, C.-C., Hillesland, K. L., Waite, A. & Shou, W. Using artificial systems to explore the ecology and evolution of symbioses. Cellular and Molecular Life Sciences 68, 1353–1368 (2011).

18. Momeni, B., Xie, L., Shou, W. & Levin, B. Lotka-Volterra pairwise modeling fails to capture diverse pairwise microbial interactions. eLife 6, e25051 (2017).

19. Grilli, J., Barabás, G., Michalska-Smith, M. J. & Allesina, S. Higher-order interactions stabilize dynamics in competitive network models. Nature 548, 210–213 (2017).

20. Oña, L. et al. Obligate cross-feeding expands the metabolic niche of bacteria. Nature Ecology & Evolution 1–9 (2021).

21. Giri, S. et al. Metabolic dissimilarity determines the establishment of cross-feeding interactions in bacteria. Current Biology 31, 5547–5557.e6 (2021).

22. Shou, W., Ram, S. & Vilar, J. M. G. Synthetic cooperation in engineered yeast populations. Proceedings of the National Academy of Sciences of the United States of America 104, 1877–1882 (2007).

23. Müller, M. J. I., Neugeboren, B. I., Nelson, D. R. & Murray, A. W. Genetic drift opposes mutualism during spatial population expansion. Proceedings of the National Academy of Sciences of the United States of America 111, 1037–1042 (2014).

24. Eigen, M. & Schuster, P. The hypercycle. a principle of natural self-organization. part a: Emergence of the hypercycle. Die Naturwissenschaften 64, 541–565 (1977).

25. Maynard Smith, J. Hypercycles and the origin of life. Nature 280, 445–446 (1979).

26. Doolittle, W. F. & Inkpen, S. A. Processes and patterns of interaction as units of selection: An introduction to ITSNTS thinking. Proceedings of the National Academy of Sciences 115, 201722232 (2018).

27. Denton, J. A. & Gokhale, C. S. Synthetic mutualism and the intervention dilemma. Life 9, 15 (2019).

28. Denton, J. A. & Gokhale, C. S. Promoting Synthetic Symbiosis under Environmental Disturbances. mSystems 5, e00187–20 (2020).

29. Baker, D. M., Freeman, C. J., Wong, J. C. Y., Fogel, M. L. & Knowlton, N. Climate change promotes parasitism in a coral symbiosis. The ISME Journal 12, 921–930 (2018).

30. Borstel, R. C. v. et al. Topical Reversion at the HIS1 Locus of Saccharomyces cerevisiae * A Tale of Three Mutants. Genetics 148, 1647–1654 (1998).

31. Yoshikawa, K. et al. Comprehensive phenotypic analysis of single-gene deletion and overexpression strains of Saccharomyces cerevisiae. Yeast 28, 349–361 (2011).

32. Gokhale, C. S. & Traulsen, A. Evolutionary games in the multiverse. Proceedings of the National Academy of Sciences USA 107, 5500–5504 (2010).

33. Morris, J. J. Black Queen evolution: the role of leakiness in structuring microbial communities. Trends in Genetics 31, 475–482 (2015).

34. Kafri, M., Metzl-Raz, E., Jona, G. & Barkai, N. The Cost of Protein Production. Cell Reports 14, 22–31 (2016).

35. Zomorrodi, A. R. & Segrè, D. Genome-driven evolutionary game theory helps understand the rise of metabolic interdependencies in microbial communities. Nature Communications 8, 6449 (2017).

36. Rivett, D. W. et al. Resource-dependent attenuation of species interactions during bacterial succession. The ISME journal (2016).

37. Caswell, H. & Trevisan, M. C. Sensitivity Analysis of Periodic Matrix Models. Ecology 75, 1299–1303 (1994).

38. Park, J. S. & Wootton, J. T. Slower environmental cycles maintain greater life-history variation within populations. Ecology Letters 24, 2452–2463 (2021).

39. Daalen, S.v. & Caswell, H. Variance as a life history outcome: Sensitivity analysis of the contributions of stochasticity and heterogeneity. Ecological Modelling 417, 108856 (2020).

40. Matsumura, S. et al. Transient compartmentalization of RNA replicators prevents extinction due to parasites. Science 354, 1293–1296 (2016).

41. Sojo, V., Herschy, B., Whicher, A., Camprubí, E. & Lane, N. The Origin of Life in Alkaline Hydrothermal Vents. Astrobiology 16, 181–197 (2016-02).

42. Pearce, B. K. D., Pudritz, R. E., Semenov, D. A. & Henning, T. K. Origin of the RNA world: The fate of nucleobases in warm little ponds. Proceedings of the National Academy of Sciences 114, 11327–11332 (2017).

43. Levings, S. C. & Garrity, S. D. Diel and tidal movement of two co-occurring neritid snails; differences in grazing patterns on a tropical rocky shore. Journal of Experimental Marine Biology and Ecology 67, 261–278 (1983).

44. Harley, C. Tidal dynamics, topographic orientation, and temperature-mediated mass mortalities on rocky shores. Marine Ecology Progress Series 371, 37–46 (2008).

45. Wasserman, B. A. et al. Ecosystem size shapes antipredator trait evolution in estuarine threespine stickleback. Oikos 129, 1795–1806 (2020).

46. Kaiser, T. S., Neumann, D. & Heckel, D. G. Timing the tides: Genetic control of diurnal and lunar emergence times is correlated in the marine midge clunio marinus. BMC Genetics 12, 49 (2011).

47. Hall, A. R. & Colegrave, N. How does resource supply affect evolutionary diversification? Proceedings of the Royal Society B: Biological Sciences 274, 73–78 (2007).

48. Hale, K. R. S., Valdovinos, F. S. & Martinez, N. D. Mutualism increases diversity, stability, and function of multiplex networks that integrate pollinators into food webs. Nature Communications 11, 2182 (2020).

49. Tonkin, J. D., Bogan, M. T., Bonada, N., Rios-Touma, B. & Lytle, D. A. Seasonality and predictability shape temporal species diversity. Ecology 98, 1201–1216 (2017).

50. Williams, C. M. et al. Understanding Evolutionary Impacts of Seasonality: An Introduction to the Symposium. Integrative and Comparative Biology 57, 921–933 (2017).

51. Yang, T. et al. Resource availability modulates biodiversity-invasion relationships by altering competitive interactions. Environmental Microbiology 19, 2984–2991 (2017).

52. Hachuy-Filho, L., Ballarin, C. S. & Amorim, F. W. Changes in plant community structure and decrease in floral resource availability lead to a high temporal beta-diversity of plant - bee interactions. Arthropod - Plant Interactions 14, 571–583 (2020).

53. Carpenter, S. R. Phosphorus control is critical to mitigating eutrophication. Proceedings of the National Academy of Sciences 105, 11039–11040 (2008).

54. Romero, E. et al. Large-scale patterns of river inputs in southwestern Europe: seasonal and interannual variations and potential eutrophication effects at the coastal zone. Biogeochemistry 113, 481–505 (2013).

55. Frumin, G. T. & Gildeeva, I. M. Eutrophication of water bodies — A global environmental problem. Russian Journal of General Chemistry 84, 2483–2488 (2014).

56. Dobson, A. P. Yellowstone Wolves and the Forces That Structure Natural Systems. PLoS Biology 12, e1002025 (2014-12).

57. Etienne, R., Wertheim, B., Hemerik, L., Schneider, P. & Powell, J. The interaction between dispersal, the Allee effect and scramble competition affects population dynamics. Ecological Modelling 148, 153–168 (2002).

58. Kaul, R. B., Kramer, A. M., Dobbs, F. C. & Drake, J. M. Experimental demonstration of an Allee effect in microbial populations. Biology Letters 12, 20160070 (2016).

59. Qin, L., Zhang, F., Wang, W. & Song, W. Interaction between Allee effects caused by organism-environment feedback and by other ecological mechanisms. PLoS ONE 12, e0174141 (2017).

60. Lindenmayer, D., Messier, C. & Sato, C. Avoiding ecosystem collapse in managed forest ecosystems. Frontiers in Ecology and the Environment 14, 561–568 (2016).

61. Dakos, V. et al. Ecosystem tipping points in an evolving world. Nature Ecology & Evolution 3, 355–362 (2019).

62. Pedraza, P. C. C., Matthews, B., Meester, L.d. & Dakos, V. Adaptive Evolution Can Both Prevent Ecosystem Collapse and Delay Ecosystem Recovery. The American Naturalist 198, E185–E197 (2021).

63. Czuppon, P. & Gokhale, C. S. Disentangling eco-evolutionary effects on trait fixation. Theoretical Population Biology 124, 93–107 (2018).

64. Doulcier, G., Lambert, A., De Monte, S. & Rainey, P. B. Eco-evolutionary dynamics of nested darwinian populations and the emergence of community-level heredity. eLife 9, e53433 (2019).

65. Williams, R. J. & Martinez, N. D. Simple rules yield complex food webs. Nature 404, 180–183 (2000).

66. Ings, T. C. et al. Review: Ecological networks – beyond food webs. Journal of Animal Ecology 78, 253–269 (2009-01).

67. Poisot, T. & Gravel, D. When is an ecological network complex? Connectance drives degree distribution and emerging network properties. PeerJ 2, e251 (2014).

68. Lorenzo, V. d. Systems biology approaches to bioremediation. Current Opinion in Biotechnology 19, 579–589 (2008).

69. Zomorrodi, A. R. & Segrè, D. Synthetic Ecology of Microbes: Mathematical Models and Applications. Journal of Molecular Biology 428, 837 –861 (2016).

70. Solé, R. V. Bioengineering the biosphere? Ecological Complexity 22, 40–49 (2015).

71. Santomartino, R., Zea, L. & Cockell, C. S. The smallest space miners: principles of space biomining. Extremophiles 26, 7 (2022).

72. Zhou, K., Qiao, K., Edgar, S. & Stephanopoulos, G. Distributing a metabolic pathway among a microbial consortium enhances production of natural products. Nature Biotechnology 33, 377–383 (2015).

73. Xie, L. & Shou, W. Steering ecological-evolutionary dynamics to improve artificial selection of microbial communities. Nature Communications 12, 6799 (2021).

74. Doebeli, M., Ispolatov, Y. & Simon, B. Towards a mechanistic foundation of evolutionary theory. eLife 6, e23804 (2017).

